# Saccades phase-locked to alpha oscillations in the occipital and medial temporal lobe enhance memory encoding

**DOI:** 10.1101/158758

**Authors:** Tobias Staudigl, Elisabeth Hartl, Soheyl Noachtar, Christian F. Doeller, Ole Jensen

## Abstract

Efficient sampling of visual information requires a coordination of eye movements and ongoing brain oscillations. Using intracranial and MEG recordings, we show that saccades are locked to the phase of visual alpha oscillations, and that this coordination supports mnemonic encoding of visual scenes. Furthermore, parahippocampal and retrosplenial cortex involvement in this coordination reflects effective vision-to-memory mapping, highlighting the importance of neural oscillations for the interaction between visual and memory domains.

## Introduction

Sampling of visual information has been shown to be rhythmic rather than continuous (1-3). In particular brain rhythms clocked by oscillations in the alpha (7-14 Hz) range (4) constrains visual sampling: EEG/MEG studies in humans have shown that the trial-by-trial fluctuations in near-threshold visual perception performance depends on the phase of alpha oscillations prior to stimulus presentation(5, 6). Saccadic eye movements overtly sample visual scenes. Here we ask how brain oscillations and saccades are coordinated, in order to allow visual information to be encoded in memory areas.

We addressed this question by tracking saccadic eye positions in separate memory experiments involving MEG in healthy adults and intracranial recordings in epileptic patients, respectively (Fig. 1). Subjects were asked to remember images of visual scenes and we later probed their memory.

**Fig 1.**
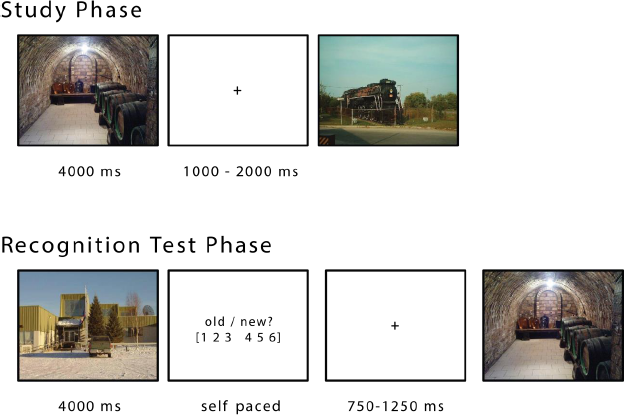
Procedure. During the study phase, participants viewed natural scenes (free viewing), indicating whether the depicted scene was indoors or outdoors (shallow encoding task). After a short distracter task (~6 min), all the images from the study phase were presented again, randomly interleaved with new pictures. Participants were prompted to judge whether a given image was new or old on a six-point scale.

The phase-locking (7) between pre-saccadic brain oscillations in relation to saccade onset was contrasted between later remembered and later forgotten images. Building on prior evidence on the cortical origins of alpha activity underlying visual information sampling (8, 9), we hypothesized that higher phase-locking in occipital lobe would result in enhanced memory performance. MEG and intracranial data both showed that eye movements are locked to the phase of alpha oscillations prior to a saccade. Importantly, this coordination was predictive of successful memory encoding.

## Results

In order to investigate the temporal coordination of saccades and brain oscillations, the time-frequency representations of phase and power of the MEG and intracranial data were aligned to saccade onsets. Accordingly, high pre-saccadic phase-locking would demonstrate an effective coordination of saccades in relation to brain oscillations. The intracranial data recorded from three patients with occipital depth electrodes (Fig. 2a) revealed a significantly higher phase-locking for later remembered as compared to later forgotten items in the alpha band (12 - 14 Hz, Cluster-randomization: p < .005, controlling for multiple comparisons over time and frequency, 2-sided test, fixed-effects statistics; Fig. 2b), prior to saccade onset.

**Fig 2.**
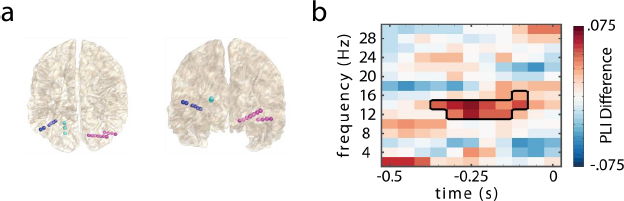
Pre-saccadic phase-locking in occipital leads of depth electrodes. (a) Electrode locations of depth electrodes in 3 epilepsy patients (color-coded). (b) Phase-locking difference (later remembered – later forgotten) in the occipital leads of the depth electrodes prior to saccade onset (t = 0 s). Significantly higher phase-locking in later remembered vs. later forgotten items; cluster highlighted by black box (p < .005, 2-sided test, fixed-effects statistics, 15 leads in bipolar montage).

The intracranial results then guided the analyses in the group level study by confining the frequency of interest to 12 - 14 Hz, where MEG data here is presented from 22 healthy participants performing the memory task. A cluster-based permutation test revealed a significant difference in pre-saccade phase-locking between later remembered and forgotten images in the alpha band (12 - 14 Hz), in a time window between −350 - −150 ms prior to saccade onset (Cluster-randomization: p < .05, controlling for multiple comparisons over time and space, 2-sided test, Fig. 3a, suppl. Fig. 1). In the posterior sensors forming a cluster, the difference was most pronounced ~250 ms prior to saccade onset (Fig. 3a).

**Fig 3.**
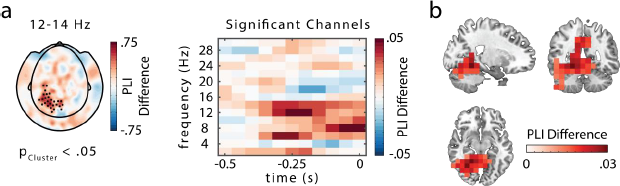
Pre-saccadic phase-locking in the MEG data. (a) MEG sensor (planar gradients) analysis showssignificantly higher phase-locking (PLI) for later remembered than forgotten items at 12 – 14 Hz, based on intracranial results (in the −0.35 – −0.15 s interval prior to saccade onset (p < .05, 2-sided test, significant channels highlighted). The time-frequency representation of the phase-locking difference averaged across highlighted sensors shows the peak of the difference to be at 250 ms prior to saccade onset (t = 0 s). (b) Phase-locking analysis (PLI) at source-level using a DICS beamformer approach. Cluster-based permutation statistics indicated a significant difference between later remembered and forgotten trials (p < .05, 2-sided test). While the maximum difference was in parahippocampal areas, the source extended to visual, parietal and temporal areas.

Analyzing the phase-locking for later remembered and forgotten pictures separately, suggested an existence of a preferred alpha phase for later remembered, but not later forgotten items (suppl. Fig. 2). Additional analyses of pre-saccadic spectral power indicated that the phase-locking results were not biased by spectral power (suppl. Fig. 3).

In order to identify the sources of the effects, we computed phase-locking in the alpha band for virtual sensors, applying a DICS beamformer(10). Cluster-based permutation statistics at the source-level yielded a significantly higher phase-locking index for later remembered than forgotten items (p < .05, 2-sided test; Fig. 3b). The cluster spanned from visual to parietal and temporal areas, extending into the cerebellum (Fig. 3b). The largest differences were found in parahippocampal gyrus and retrosplenial cortex that have been shown to support the encoding of visual scenes, extending into posterior hippocampus. The MEG source localization is supported by intracranial data from parahippocampal depth electrodes in the three patients, showing significantly higher phase-locking for later remembered versus later forgotten items in the alpha range (8 - 10 Hz, p cluster < .05; 2-sided test, fixed-effects statistics; suppl. Fig. 4).

## Discussion

In two independent data sets, we provide novel evidence for a functionally relevant coordination of saccadic eye-movements and brain activity. Both the intracranial and MEG data show that retinal inputs are temporally aligned to a preferential alpha phase. Importantly, this coordination resulted in enhanced memory encoding, suggesting a mechanistic role for alpha oscillations in coordinating the encoding of visual information. Furthermore, our results point to an active involvement of task-relevant brain areas in this coordination: MEG and intracranial data yielded occipital, parahippocampal gyrus and retrosplenial cortex as sources of the coordination of saccades and alpha phase, which have been shown to support the encoding of visual scenes (11-13). The engagement of scene-selective areas may reflect effective vision to memory mapping along posterior cortex (14).

Our findings are in line with work from the 1960s (15) suggesting a relationship between alpha oscillations and saccades; however, this effect was not related to perception and memory. They also support the notion of a preferred alpha phase for the execution of eye movements(16), by suggesting that during optimal information encoding, the execution of saccades is on hold until the end of an alpha duty cycle. We propose that effective coordination of saccades and brain oscillations allows for optimizing the speed of processing in the visual system (17).

The increase in memory encoding with saccades locked to alpha phase might be supported by anticipatory attentional deployment (18). The fact that the phase-locking difference was found prior to saccade onset might suggest planning of the upcoming to-be-attended location(19) resulting in a stronger locking of saccades to the phase of the alpha oscillation and ultimately improved memory encoding. The present results highlight the necessity for a coordination of alpha oscillations and eye movements for optimal memory encoding. Efficiently sampled visual information could then be integrated by the hippocampal memory system. A recent non-human primate study demonstrated that saccades were aligned to hippocampal ~10 Hz oscillations (20). Future studies should explore interregional synchronization in relation to oculomotor behavior, visual information sampling and memory.

## Materials and Methods

### Participants

For the MEG part, 36 young healthy adults were included in the study. Initially, 48 participants were recruited; however, 12 dropped out due to not completing the study (7 participants), excessive movement artifacts (2 participants) and technical problems during the recordings (3 participants). All participants gave written informed consent before the start of experiment in accordance with the Declaration of Helsinki. The study was approved by the local ethics committee (commission for human related research CMO-2014/288 region Arnhem/Nijmegen NL). The 36 participants included in this study (24 females; mean age 23.1 years, range 18-30 years; 35 right handed) reported no history of neurological and/or psychiatric disorders and had normal or corrected-to-normal vision.

Additionally, three male patients (age range 30-60 years) with a history of drug resistant epilepsy were recruited from the Epilepsy Center, Department of Neurology, University of Munich, Germany. The patients, who volunteered to participate in the study, had depth electrodes implanted for diagnostic reasons. The patients gave written informed consent. The study was approved by the ethics committee of the University of Munich.

### Design, Procedure and Materials

The study design comprised an MEG and an fMRI (not reported here) session. Session order was counterbalanced across participants. For each session, three stimulus sets of 100 photographs each were constructed. Half of the pictures depicted indoor scenes, the other half outdoor scenes (exemplary scenes are shown in Fig. 1). Two sets were presented during encoding, one during test as foils. Assignment of set to encoding or test was counterbalanced across participants. Nine additional scenes were presented during a short practice session before encoding and test in order to explain the task.

Fig. 1 illustrates the experimental procedure. At study, the pictures were presented for 4 s in random order with the constraint that no more than four scenes of the same type (indoor / outdoor) were shown consecutively. The participants were instructed to judge whether the depicted scene was indoors or outdoors by button press. This encoding task was chosen to ensure attention to each scene and promote encoding of the images. Participants freely viewed the figures; i.e. they were not expected to fixate. A fixation cross with variable duration (1 - 2 s) followed each scene.

The study phase was followed by a distracter phase during which the participants solved simple mathematical problems for ~ 1 min, ~5 min of fixation to different locations on the screen used to evaluate eye tracker accuracy, ~1 min of eyes open and ~1 min of eyes closed. The distracter phase prevented participants from covert rehearsing. After the distractor, period followed the memory recall. Participants were instructed about the memory test before the start of the experiment. At test, the 200 pictures from the study phase and 100 new pictures (foils) were presented for 4 s. The presentation order was randomized, with the constraint that no more than four scenes of the same type (old / new) were shown consecutively. After each scene, participants were prompted to indicate their confidence on whether the item was old or new using a six-point response scale, ranging from 'very sure old’ (1) to 'very sure new’ (6). This picture of the rating scale remained until the participants responded. Before the next scene, a fixation-cross with variable duration (750 - 1250 ms) was presented. The procedure for the patients with intracranial electrodes deviated slightly (see below).

### MEG Acquisition and Preprocessing

MEG was recorded using a 275 whole-brain axial gradiometer system (VSM MedTech/CTF MEG, Coquitlam, Canada) installed in a magnetically shielded room. The data were sampled at 1200 Hz following a low-pass anti-aliasing filter with a cutoff at 300 Hz. Additionally, horizontal and vertical electrooculograms were recorded from bipolar Ag/AgCl electrodes (<10kQ impedance) placed below and above the left eye and at the bilateral outer canthi. To track the position of the head during MEG recording, we used 3 head coils placed at anatomical landmarks (nasion and both ear canals). Using a real-time head localizer(21), the position of the head relative to the MEG helmet was tracked. The participants' nasion, left and right ear canal, and head shape were digitized with a Polhemus 3Space Fasttrack.

Preprocessing of the data was done using the Fieldtrip toolbox(22). Data were divided into single trials, with epochs ranging from, 0 to 4 s after picture onset. Trials were corrected for cardiac artifacts using Independent Component Analysis (ICA) and sorted according to the behavioral performance of each participant's confidence judgments during the recognition test phase. Trials that were confidently judged as old (responses 1, 2, and 3) constituted hits, the remaining trials were classified as misses.

### Eye Tracking Acquisition, Analyses and Trial Definition

An Eyelink 1000 (SR Research) eyetracker was used to monitor the horizontal and vertical movements of the participants' left eye. Before recording, the eye tracker was calibrated by collecting gaze fixation samples from known target points to map raw eye data on screen coordinates. Participants fixated on nine dots sequentially on a 3 by 3 grid. After the calibration run, a validation run was performed during which the difference between current gaze fixations and fixations during the calibration were obtained. The calibration was only accepted, if this difference was smaller than 1 degree visual angle.

Eye tracking and MEG data were simultaneously recorded and analyzed using the Fieldtrip toolbox. Vertical and horizontal eye movements were transformed into velocities. Velocities exceeding a certain threshold (velocity > 6 x the standard deviation of the velocity distribution, duration > 12 ms, see Engbert and Kliegl (23)) were defined as saccades. Saccade onsets during stimulus presentations defined the events of interests (trials). To avoid potential artifacts from other eye movements, only events that were free of saccades and blinks in a 0.5 s interval prior to saccade onset were included. After excluding all participants that had less than 30 remaining trials, 22 participant remained for further analyses.

### Phase and power analysis

The time-frequency representations of phase and power the data were computed by a sliding time window approached with a window length of 0.5 s in steps of 50 ms across the data. After multiplying a hanning taper to each window, the Fourier transformation was calculated for frequencies between 2 and 30 Hz in steps of 2 Hz. Synthetic planar gradient representations were approximated by relating the field at each sensor with its neighbors’(24). On each of the resulting two orthogonal gradients, Fourier coefficients were normalized by their amplitude and the phase-locking index (7) (PLI) was calculated, by extracting the length of the resulting vector after averaging the phase angles:

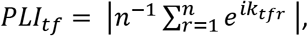

with n = number of trials, and *e^ik^* equals the complex polar representation of phase angle k in trial r, for time-frequency point tf.

This was done for later forgotten and later remembered trials, respectively. The PLI quantifies the consistency of phases across trials at each given time-frequency point. To control for a bias in PLI due to different trial numbers in conditions, a sample of trials from the condition with the larger number of trials was randomly drawn, with the number of trials in this sample being equal to the number of trials in the condition with less trials. The PLI for this sample was computed. After repeating this procedure 1000 times, PLI values were averaged. This average reflects an unbiased estimate of the PLI for all trials in the respective condition. After this step, the two planar gradients were combined.

To identify potential confounds due to differences in spectral power, power was calculated on synthetic planar gradients, using the same approach as outlined above. Instead of computing the PLI, power values were calculated from the Fourier coefficients (amplitude squared).

### Source level analyses

To identify PLI differences in source space, a virtual sensor approach applying frequency domain adaptive spatial filtering (DICS beamformer (10)) was implemented. This algorithm constructs a spatial filter for each specified location (each grid point; 10mm^3^ grid). The cross-spectral density for the construction of the spatial filter was calculated for the frequency and time window of interest (12 - 14 Hz; −0.35 - −0.15 ms) by means of Fourier transformation and hanning taper (see above), for all trials (common filter approach).

Individual structural MR images, acquired on a 3T Siemens Magnetom Prisma MRI system (Siemens, Erlangen, Germany) were aligned to the MEG coordinate system, utilizing the fiducials (nasion, left and right preauricular points) and individual head-shapes recorded after the experiment. A realistic singleshell brain model (25) was constructed for each participant, based on the structural MRIs. The forward model for each participant was created using a common dipole grid (10mm^3^ grid) of the grey matter (derived from the anatomical automatic labeling atlas (26)) volume in MNI space warped onto each participant's anatomy. The Fourier data was projected into source space by multiplying it with the spatial accordant filters allowing for the phase to be estimated. The PLI was computed on the two orientations of the source model, and later averaged, for later remembered and later forgotten items, respectively.

### Statistics

Statistics followed a two-step approach: First, differences in the intracranial data's phase-locking (later remembered versus later forgotten items) were evaluated in a fixed-effect manner, by concatenating all electrodes from all patients. Cluster-based nonparametric permutation statistics(27) identified continuous time-frequency clusters with significant differences between later remembered and later forgotten PLI, while controlling for multiple comparisons over time and frequency. Only the cluster with the largest summed value was considered and tested against the permutation distribution. The null-hypothesis that later forgotten and later remembered trials showed no difference in PLI was rejected at an alpha-level of 0.05 (two-tailed).

Second, statistical quantification of the MEG sensor level data was performed by a cluster-based nonparametric permutation approach (27), identifying clusters of activity on the basis of rejecting the null hypothesis while controlling for multiple comparisons over sensors and time-points. The frequency range (12 - 14 Hz) for the sensor level statistics was restricted to the outcome of the intracranial data analyses. For each sensor, a test statistic was calculated, based on a paired samples t-test comparing the PLI for later remembered vs. later forgotten trials. Sensors showing a significant effect (p < 0.05, two-sided t-test), were clustered based on spatial adjacency, with a minimum of 2 adjacent sensors required for forming a cluster. T-statistics were summed in each cluster. Again, only the cluster with the largest summed value was considered and tested against the permutation distribution. The null-hypothesis that later forgotten and later remembered trials showed no difference in PLI was rejected at an alpha-level of 0. 05 (two-tailed).

Statistical quantification of the source level data was also performed by a cluster-based nonparametric permutation approach, now considering the clustering in voxel space. The time-frequency range for the source level statistics was defined by the outcome of the sensor level statistics, alpha level was set to 0.05 (two-tailed). Cluster-based nonparametric permutation statistics (27) identified continuous spatial clusters with significant differences between later remembered and later forgotten PLI, while controlling for multiple comparisons over voxels. Only the cluster with the largest summed value was considered and tested against the permutation distribution. The null-hypothesis that later forgotten and later remembered trials showed no difference in PLI was rejected at an alpha-level of 0.05 (two-tailed).

Condition specific PLIs for later remembered and later forgotten trials, separately, (see suppl. Fig. 2) at the time and frequency of interest (12 - 14 Hz, −0.35 - −0.15 ms) were statistically quantified by comparing them to a distribution of surrogate PLI values. The surrogate PLI distribution was constructed for each participant and condition by shifting the data points in each condition's trial circularly along the time axis with a random lag, for each channel. PLI values were computed as explained above, for 1000 random shifts. Subsequently, 10000 surrogate grand averages were constructed by randomly drawing one PLI value from each subject's surrogate distribution for each surrogate grand average. Conditions specific PLI grand averages were compared to these 10000 surrogate grand averages on each sensor, and considered to be significant if they were larger (or smaller) than the 97.5 % (or 2.5 %) of the values in the surrogate grand average values (two sided test).

### Intracranial Data

Three male patients (age range 30-60 years) with occipital depth electrodes were included in the study. The patients had a history of drug resistant focal epilepsy and were implanted for diagnostic reasons. Recordings were performed at the Epilepsy Center, Department of Neurology, University of Munich, Germany. The patients gave written informed consent. The procedure and design of the study was identical to the MEG procedure and design (see above), with the exception that only 100 pictures were presented during study and 200 scenes (100 old and 100 new) were presented during the memory test. This was done to compensate for inferior memory performance in a clinical setting.

Patient 1 had ten depth electrodes implanted, covering bilateral temporal, parietal and frontal regions and left occipital regions. Patient 2 had ten depth electrodes implanted, covering right temporal, parietal and occipital regions. Patient 3 had eleven depth electrodes implanted, covering left frontal, temporal, parietal and occipital regions. The locations of the electrodes were determined using coregistered preoperative MRIs and postoperative CTs. Electrode locations were converted to MNI coordinates. Intracranial EEG was recorded from Spencer depth electrodes (Ad-Tech Medical Instrument Corporation, Racine, USA) with 4-12 contacts each, 5mm apart. Data was recorded using XLTEK Neuroworks software (Natus Medical Incorporated, San Carlos, USA) and an XLTEK EMU128FS amplifier, with voltages referenced to a parietal electrode site (1000 Hz sampling rate). All electrodes that either were identified as located in the seizure onset zone or showed interictal spiking activity were excluded from analyses.

Data was re-referenced offline to each contact's neighboring contact (bipolar montage). All bipolar electrodes with both contacts in the occipital cortex were included in the analyses. Additionally, horizontal and vertical eye movements were recorded from bipolar Ag/AgCl electrodes (<10kQ impedance) placed below and above the left eye and at the bilateral outer canthi.

Study phase data were epoched into single trials, with epochs ranging from, 0 to 4 s after picture onset. Saccade onsets were extracted from EOG recordings using the method described above (see Eye Tracking Acquisition, Analyses and Trial Definition). All trials were visually inspected for artifacts (e.g. epileptoform spikes). Contaminated trials were excluded from the analyses. The encoding trials were sorted according to each participant's confidence judgments during the test phase. Trials including old pictures that were judged as old (responses 1, 2, and 3) constituted hits; the remaining trials including old pictures were classified as misses. Time-frequency analyses, PLI and statistics were computed as described above

## Acknowledgments

We thank all of the participants, in particular the patients, for taking part in the study, and the technical staff at the Epilepsy Center Munich for their support. This project has received funding from the European Union's Horizon 2020 research and innovation programme under grant agreement No 661373. CFD's research is funded by the Kavli Foundation, the Centre of Excellence scheme of the Research Council of Norway - Centre for Biology of Memory and Centre for Neural Computation, The Egil and Pauline Braathen and Fred Kavli Centre for Cortical Microcircuits, the National Infrastructure scheme of the Research Council of Norway - NORBRAIN, the Netherlands Organisation for Scientific Research (NWO-Vidi 452-12-009; NWO-Gravitation 024-001-006; NWO-MaGW 406-14-114; NWO-MaGW 406-15-291) and the European Research Council (ERC-StG RECONTEXT 261177; ERC-CoG GEOCOG 724836).

